# Survey of severe fever with thrombocytopenia syndrome virus covert infection for healthy people in Henan Province, China

**DOI:** 10.1101/550426

**Authors:** Yanhua Du, Ningning Cheng, Yi Li, Haifeng Wang, Aiguo You, Jia Su, Yifei Nie, Hongxia Ma, Bianli Xu, Xueyong Huang

**Author notes:** Corresponding author (Bian-li Xu) (Xue-yong Huang). These first authors contributed equally to this article (Yanhua Du, Ningning Cheng).

## Abstract

**Background:** Severe fever with thrombocytopenia syndrome (SFTS) is a severe emerging disease, and its incidence has been increasing in recent years. A cross-sectional study was conducted for healthy people in high SFTS endemic areas of Henan province in 2016.

**Methods:** This study used stratified random sampling method and finally 14 natural villages were selected as the investigation site. The questionnaire survey and serum collection were carried out from April to May in 2016. All the serum samples were detected by SFTSV IgM and IgG antibodies by ELISA. Only positive samples of SFTSV IgM antibody need be tested SFTSV RNA and virus cultured. A month after the specimen collection, all persons positive for IgM antibody were followed up one by one to confirm whether he or she was recessive infection.

**Results:** 1463 healthy persons were investigated in total. The average seropositive rates of SFTS virus specific IgG and IgM antibodies were 10.46% (153/1463) and 0.82% (12/1463), respectively. IgM antibody was positive in 12 individuals, and 6 of them SFTS virus RNA were also detected. In addition, 5 SFTS virus were isolated from the 6 SFTS virus RNA (+) individuals. By genomic sequencing and phylogenetic analysis, 5 new isolates from healthy crowd all belonged to groups A, which were similar to other isolates from SFTS patients. After one month, all people with SFTS virus IgM antibody had no illness or symptoms.

**Conclusions:** This study confirmed there is SFTS recessive infection in human, and it is the first report about SFTS virus isolation in healthy people.

**Author Summary:** Severe fever with thrombocytopenia syndrome (SFTS), a severe emerging infectious disease, was discovered in rural areas of China. The first SFTS case was found in Henan province, which has had the largest number of SFTS cases in China every year since the disease was discovered. However, as a high incidence area of SFTS in Henan province, the serum prevalence rate of the virus in healthy people is still not clear. Therefore, a cross-sectional survey was performed in high endemic areas and epidemic seasons in 2016. The results showed that the level of specific SFTS seroprevlence was relatively higher and possibility increasing. SFTS RNA were positive and SFTS virus was isolated from the specimens of healthy people. This study confirmed there is SFTS recessive infection in human, and it is the first report about SFTS virus isolation in healthy people.

## Introduction

Severe fever with thrombocytopenia syndrome (SFTS) is a kind of emerging infectious disease, which appeared in eastern China since 2007[1,2]. The major clinical syndromes include fever, thrombocytopenia, leukocytopenia, gastrointestinal symptoms, neurological symptoms, bleeding tendency, as well as less specific clinical manifestations[3,4]. Until 2009, the severe fever with thrombocytopenia syndrome virus (SFTS virus), a novel virus, was first identified from a patient located in Xinyang, Henan, China[5]. The SFTS patients had been reported in almost 25 provinces of China and other countries such as Japan and Korea[6,7]. SFTS Virus was listed as concerned special pathogens in 2016 and 2017 by World Health Organization (WHO).

In recent years, there were various epidemiologic studies for seroprevalence of SFTS virus among healthy humans in different sampling method and size, region, time, and so on. A recent systematic review and meta-analysis showed the overall pooled seroprevalence of SFTS virus was 4.3%,ranged from 0.23% to 9.17%[8]. The SFTSV infections rates of healthy humans varied in different regions of China, which is likely caused by limiting number of study and sample size in endemic regions. In China, more than 85% of SFTS cases lived in rural regions of seven provinces, and the number of infected cases in Henan province always took the first place since the novel SFTS virus was discovered[9,10]. Therefore, it is important to disclose the actual seroprevlence of SFTS virus and to further estimate for healthy individuals in high endemic areas.

## Methods

### Study Design

It was reported that SFTS virus was discovered in Henan province, and there is the most SFTS cases amount in China, as well as more than 95% of SFTS cases in Henan Province reported came from Xinyang City which is a high endemic areas[11]. A cross-sectional investigation was conducted in Xinyang city by random cluster sampling. The city is divided into 10 administrative counties/districts. First, one county (Xin) and one district (Pingqiao) were selected; second, two towns (Balifan, Pengjiawan) were selected from Xin county and Pingqiao district, and then, fourteen natural villages were selected from Balifan and Pengjiawan town, respectively. The healthy people in this study were from fourteen natural villages that had SFTS reported cases before.

The survey subjects were selected with strict inclusion criteria and exclusion criteria, who must live in the area for more than 1 year, ages 2 years and older, and were not SFTS patients in the past. The previous studies have shown SFTS virus seroprevalence was 7.2%-10.5% in healthy persons who reported no symptoms associated with SFTS[12]. The estimation of sample size depends on 7% incidence, 80% power and two tailed 5% level of significance in this study, the minimal sample size was calculated to be 1276.

### Field Investigation and Sample Collection

From April to May 2016, the survey subjects were investigated in fourteen selected natural villages. A questionnaire was used to collect information on name, gender, age, occupation, home address, whether had fever two weeks or SFTS patients in the past, and et al. Serum samples were collected from survey subjects who meet the inclusion criteria. All samples were transported frozen to the pathogen laboratory of the Henan Center for Disease Control and Prevention (Henan CDC).

### Indirect Enzyme-linked Immunosorbent Assays (ELISA)

For the indirect IgG ELISA, 96-well plates were coated with 50μg recombinant nucleoprotein of SFTS virus per well overnight at 4°C. Recombinant nucleoprotein of SFTS virus was expressed and purified as our previously described[13]. Plates were washed three times with wash buffer (0.01M PBS, 0.05% Tween 20). Serum samples were diluted at 1:400 in 5% nonfat milk with PBS-T. 50μL of diluted serum samples added to each well and incubated at 37°C for 1 hour, and then washed plates with wash buffer. Plates were incubated with 1:30,000 diluted horseradish-peroxidase-conjugated goat anti-human IgG (American Qualex, California, USA) at 37°C for 1 hour. After washed, plates with 100μL H2O2-ABTS substrate (Kirkegaatrd&perry, Gaithersburg, MD) in each well were incubated for 30 min at 37°C. Absorbance was read at 405nm (including negative and blank controls). The test results were determined to be valid if the criteria for the positive control and the negative controls were fulfilled. Calculation of the cut-off value was the mean absorbance value for negative controls times 2.1; if the mean absorbance value for negative controls was lower than 0.05, then the value 0.05 was used. The positive/negative determination of samples was performed using the cut-off value.

The procedure for indirect IgM ELISA was similar to indirect IgG ELISA. The difference was that the detection of bound IgM was done with 1:10,000 diluted horseradish-peroxidase-conjugated goat anti-human IgM(American Qualex, Califonia, USA).

### Detection of SFTS virus by real-time RT-PCR

Total RNA was extracted from SFTS virus IgM positive samples using a QIAamp viral RNA mini Kit (Qiagen, Germany), following the manufacturer’s instructions. The real-time RT-PCR assay using PCR Diagnostic Kit for SFTS virus RNA (BGI-GBI, China) was performed as previously described[14]. Data were analyzed using the software supplied by the manufacturer.

### Virus Isolation and Sequencing

All the SFTS virus RT-PCR positive samples were used to inoculate on Vero cells. 100μL of serum was inoculated onto Vero cell monolayers and incubated for 7-10 days at 37°C 5% CO2 in MEM/1% fetal calf serum. All cultures were monitored daily for CPE. Both virus-infected cells and negative control cells were examined by SFTS virus real-time RT-PCR. The criteria of SFTS virus isolation and identification in tissue culture are similar with other virus isolation. Virus replication cause the cytopathic effect (CPE) in cell culture passages, and virus-positive cells were identified by real-time RT-PCR. If CPE didn’t be appeared in samples culture and the titers of SFTSV RNA didn’t increased after 3 passages, these samples were judged as negative and excluded28. The whole genome sequences of SFTS virus isolated were amplified using primers described in previous studies[15]. The RT-PCR products were sent to Jinsirui Biotech Co., Ltd (Nanjing, China) for DNA sequencing using an automated ABI 3730 DNA Sequencer.

### Phylogenetic Analysis

The genomes of SFTS virus isolates were compiled using the SeqMan program in the LaserGene software package (DNAStar). The percentage similarities of nucleotide identity or amino acid identity were calculated using the ClustalX software[16]. Molecular phylogenetic analysis was conducted by using the maximum likelihood (ML) method based on the Kimura 2-parameter model in the MEGA 5 software. SFTS virus nucleotide sequences of genome segments from GenBank were analyzed, together with our newly generated genome of SFTS virus isolates in this study, and the phylogenetic trees were constructed in order to understand the evolutionary characterization of SFTS virus.

### Statistical Analysis

All epidemiologic and laboratory data were double-entered with the EpiData 3.1 software. All statistical analyses were performed using SAS v9.13 (SAS Institute Inc., Cary, NC). The significance level α was 0.05.

### Ethics Statement

This research was approved by the Institutional Review Board at the Center for Disease Control and Prevention of Henan Province (NO.2016-KY-002-02). The methods were carried out in accordance with the principles of the Declaration of Helsinki. All adult participants signed the written informed consent and agreed to use their serum samples for research. The child participants signed the written informed consent by their parent or legal guardian of participants.

## Results

### Survey Subjects and Serum samples

A total of 1463 healthy persons were surveyed in high SFTS epidemic areas, including 768 from Xin county and 695 from Pingqiao district. Among the survey subjects, 409 were males and 1054 were females, with a male–female sex ratio of 0.39. The median age of the survey subjects was 60 years (range: 2–92). Every survey subjects gave sera, and 1463 serum samples were collected in the study.

### SFTS virus IgG and IgM antibody prevalence

All serum samples were tested for SFTS virus IgG and IgM antibodies by indirect ELISA. 153 positive IgG antibodies and 12 positive IgM antibodies were tested from 1463 healthy individuals. The seropositive rate for IgG antibody was 10.46% (153/1463), and that was 0.82% (12/1463) for IgM antibody. These SFTS virus IgG seropositive rates of gender and age were not statistically different by chi square test (χ2=0.022, P=0.883; χ2=10.076, P=0.121), and the SFTS virus IgM seropositive rates of gender and age were also not statistically different (χ2=0.009, P=0.925; χ2=7.147, P=0.307) (Table 1).

**Table 1.** Demographic characteristic of SFTS virus IgG and IgM seropositive in healthy people

| Variable | NO. | Positivefor IgG<br>(ratio,positive/total) | Positive for IgM<br>(ratio,positive/total) | Constituent ratio |
| --- | --- | --- | --- | --- |
| **Gender** |  |  |  |  |
| Males | 409 | 42 (10.27%) | 4 (0.98%) | 27.96% |
| Females | 1054 | 111 (10.53%) | 8 (0.76%) | 72.04% |
| Total | 1463 |  |  |  |
| $\chi^2$ | | 0.022 | 0.009* | |
| P |  | 0.883 | 0.925 |  |
| **Age (years old)** |  |  |  |  |
| ≤20 | 36 | 2 (5.56%) | 0 (0.00%) | 2.46% |
| 21~30 | 19 | 1 (5.26%) | 0 (0.00%) | 1.30% |
| 31~40 | 59 | 6 (10.17%) | 2 (3.39%) | 4.03% |
| 41~50 | 264 | 16 (6.06%) | 3 (1.14%) | 18.05% |
| 51~60 | 390 | 48 (12.31%) | 1 (0.26%) | 26.66% |
| 61~70 | 437 | 54 (12.36%) | 4 (0.92%) | 29.87% |
| >70 | 258 | 26 (10.08%) | 2 (0.78%) | 17.63% |
| Total | 1463 | 153 (10.46%) | 12 (0.82%) |  |
| $\chi^2$ | | 10.076 | 7.147 | |
| P |  | 0.121 | 0.307 |  |

### Real-time PCR for SFTS virus on IgM seropositive samples and isolation virus

12 positive IgM antibodies were tested from 1463 healthy individuals, and 6 of them SFTS virus RNA were also detected by RT-PCR. (Table 2) 5 isolated SFTS virus culture appeared the cytopathic effect (CPE)in cell culture passages. Compared with control Vero cells, the main expressions of CPE in infected Vero cells include: decreased cell diopter, rounding reduced cells, and modified particles. (Figure 1) The titers of SFTSV RNA of 5 isolates in the culture supernatants increased obviously after 3 passages. (Table 2)

**Table 2.**
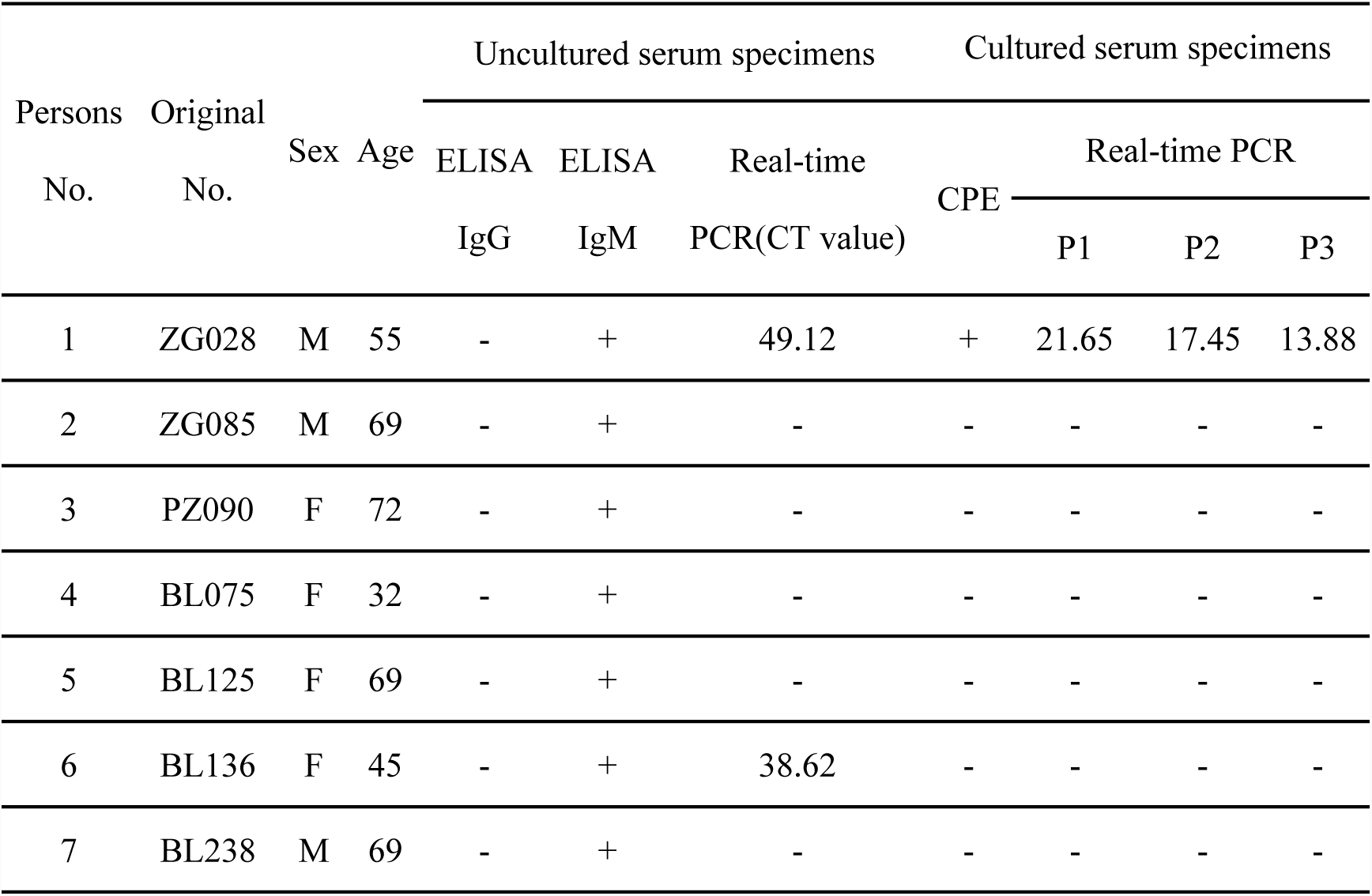
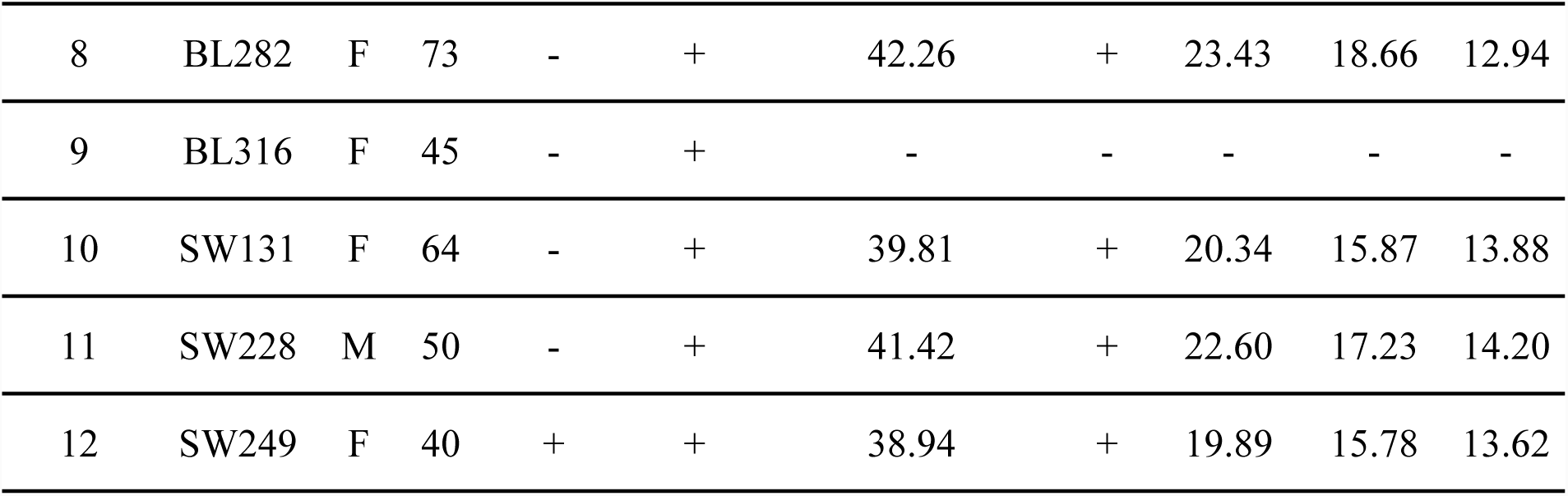
Results of ELISA, IFA and real-time RT-PCR in 12 IgM seropositive healthy persons

| Persons<br>No. | Original<br>No. | Sex | Age | Uncultured serum specimens |  |  | CPE | Cultured serum specimens |  |  |
| --- | --- | --- | --- | --- | --- | --- | --- | --- | --- | --- |
|  |  |  |  | ELISA | ELISA | Real-time |  | Real-time PCR |  |  |
|  |  |  |  | IgG | IgM | PCR(CT value) |  | P1 | P2 | P3 |
| 1 | ZG028 | M | 55 | - | + | 49.12 | + | 21.65 | 17.45 | 13.88 |
| 2 | ZG085 | M | 69 | - | + | - | - | - | - | - |
| 3 | PZ090 | F | 72 | - | + | - | - | - | - | - |
| 4 | BL075 | F | 32 | - | + | - | - | - | - | - |
| 5 | BL125 | F | 69 | - | + | - | - | - | - | - |
| 6 | BL136 | F | 45 | - | + | 38.62 | - | - | - | - |
| 7 | BL238 | M | 69 | - | + | - | - | - | - | - |
| 8 | BL282 | F | 73 | - | + | 42.26 | + | 23.43 | 18.66 | 12.94 |
| 9 | BL316 | F | 45 | - | + | - | - | - | - | - |
| 10 | SW131 | F | 64 | - | + | 39.81 | + | 20.34 | 15.87 | 13.88 |
| 11 | SW228 | M | 50 | - | + | 41.42 | + | 22.60 | 17.23 | 14.20 |
| 12 | SW249 | F | 40 | + | + | 38.94 | + | 19.89 | 15.78 | 13.62 |

**Figure 1.**
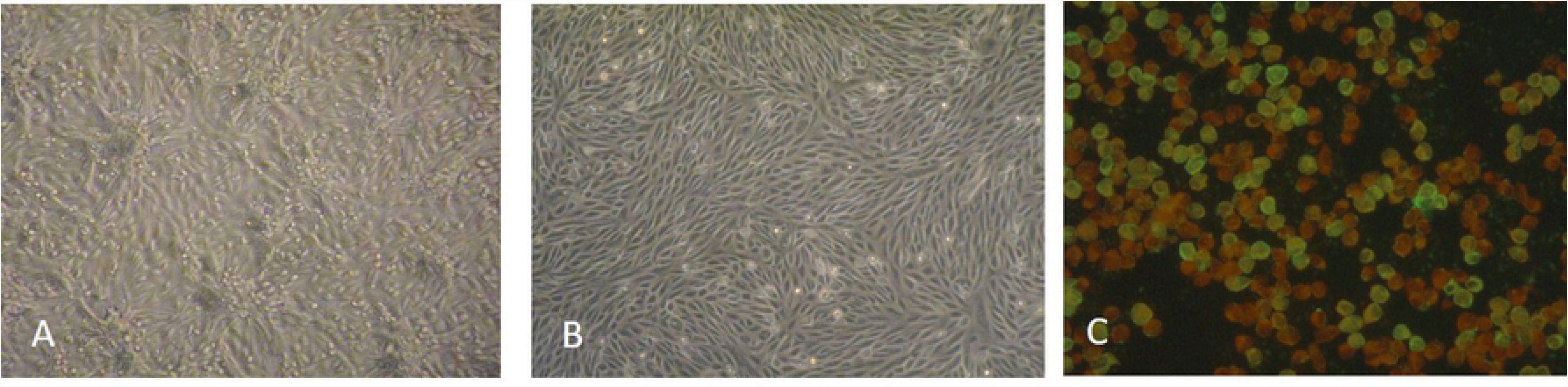
Morphologic Features of SFTSV strains isolated from healthy volunteers Panel A shows SFTS virus-induced cellular changes that are visible on light microscopy (cytopathic effect) in Vero cells 8 days after inoculation. Panel B shows mock-inoculated Vero cells. Panel C shows virus grown in Vero cells and detected on immunofluorescence assay with a serum specimen obtained from a convalescent SFTSV patient.

### Molecular Characterization of SFTS virus Strains

The complete genomes of five SFTS virus strains were successfully amplified and sequenced. The SFTS virus genomic segments (L, M and S) obtained in this study were all published in GenBank (Accession number: MF045948-MF045952, MF045955-MF045959, MF045962-MF045965 and MF045968). The nucleotide sequences of five SFTS virus isolates were closely related to each other, with 99.9% nucleotide identity for the complete L segments, 99.9% to 100% nucleotide identity for the complete M segments and the complete S segments. The identity of amino acid sequences was 99.9% to 100% in RNA-dependent RNA polymerase and Gn Gc precursor, and 100% in N protein and NSs protein, respectively.

Using the ML method in MEGA software, phylogenetic trees were constructed with the sequences of 5 isolates in this study and 30 reference SFTS virus strains from GenBank. Phylogenetic analysis of the L segment sequences revealed that the SFTS virus strains were classified into four large groups in a previous study: A, B, C and D. According to the classification, the five new isolates all belonged to groups A, and were similar to other isolates from SFTS virus patients (Figure 2).

**Figure 2.**
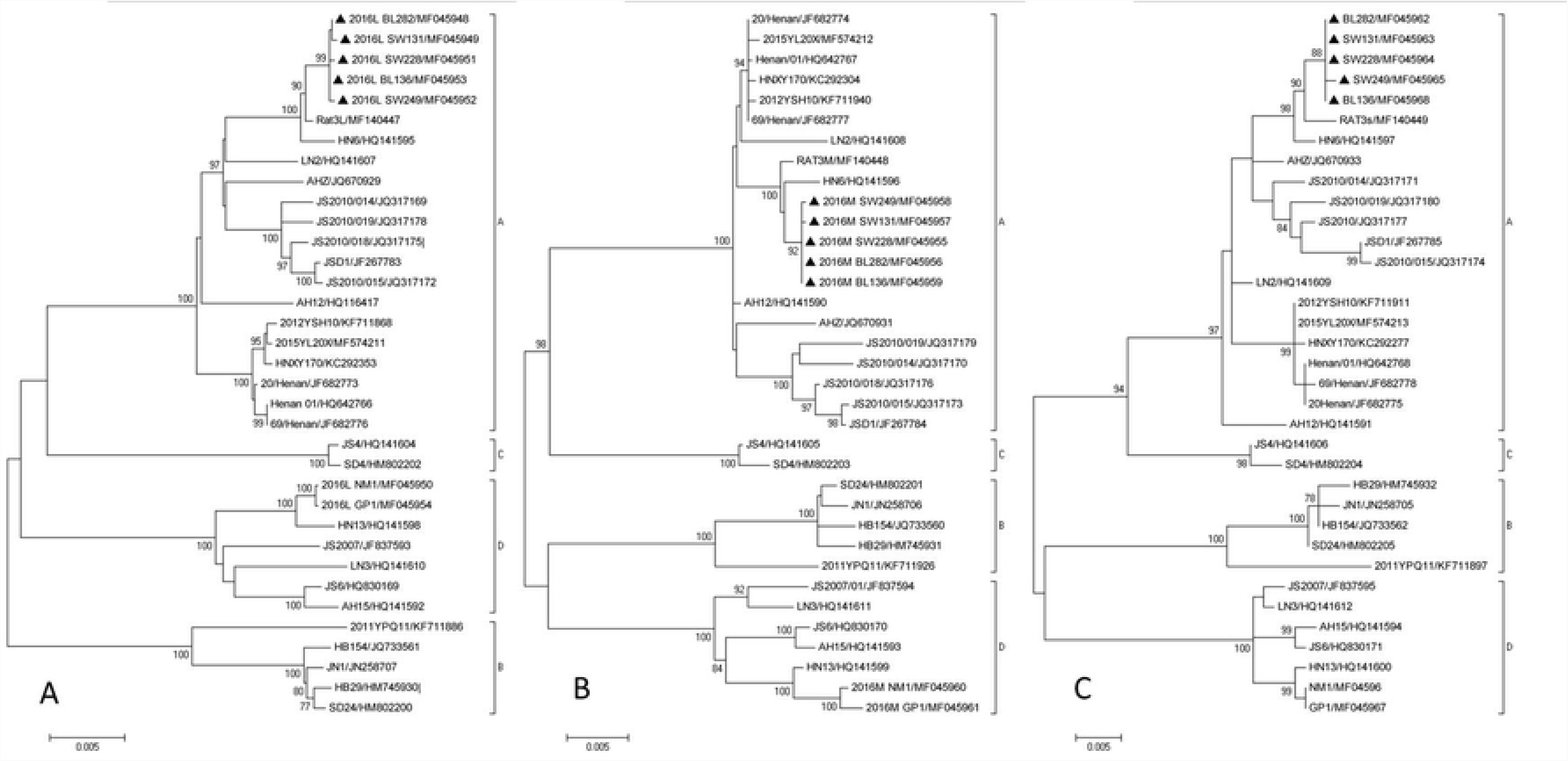
Phylogenetic analysis of SFTS virus strains isolated from healthy people from Xinyang, Henan compared with other SFTS virus. The phylogenetic tree was constructed by the maximum likelihood method with the MEGA5 software. The reliability values indicated at the branch nodes were determined using 1,000 bootstrap replications. Isolated SFTS virus strains in this study were labeled by black solid diamonds. Phylogenetic relationship of SFTS virus with other bunyaviruses, based on the complete L, M, S segment sequences, are shown in panel A, B, C, respectively. STROBE Statement—checklist of items that should be included in reports of observational studies

### Follow-up investigation

12 healthy persons with SFTS virus IgM antibody were followed up after one month, they all had no clinical symptoms of SFTS. The median age of the 12 individuals was 60 (range, 32-73) years, and the sex ratio was 1 to 2 (4 males and 8 females). The healthy persons gave serum, all serum samples were negative for SFTS virus RNA by real-time PCR, negative for SFTS virus IgM antibodies but positive for SFTS virus IgG by indirect ELISA, which showed they were recessive infection like other survey subjects.

## Discussion

Severe fever with thrombocytopenia syndrome (SFTS) is an emerging tick-borne zoonotic virus, which was first described in rural regions of China, also reported in Korea, Japan and Dubai[17-20]. Now it is an expanding global public health threat due to its increased incidence area and high mortality [21,22]. The average case fatality rate of SFTS was about 30% when this disease was firstly reported [5]. A recent report showed that an estimated 8.3% of SFTS diagnose cases were missed in high endemic areas [23]. The missed diagnosis reason is that some SFTS lab-confirmed cases lacked typical syndromes such as fever or thrombocytopenia, which means the actual incidence of SFTS may be much higher than the currently reported level. It is likely to be underestimated the status of SFTS virus in healthy people as well. Therefore, a cross-sectional study was performed to evaluate the covert infection of seroprevlence of SFTSV in Xinyang rural region.

In this study, we investigated 1463 healthy persons in 14 natural villages of Xinyang city from April to May, 2016. The average seropositive rate of SFTS virus-specific IgG antibodies was 10.46% (153/1463). To our surprise, 12 positive IgM antibodies were tested from 1463 healthy individuals, and 6 of them SFTS virus RNA were also detected by RT-PCR, as well as SFTS virus were isolated from 5 of the 6 SFTS virus RNA (+). It was first reported that SFTS virus could be isolated from healthy individuals.

In past years, seroprevalence of SFTS virus among healthy population has been investigated in various epidemiologic studies, but seroprevalence of SFTS virus among general humans in China are almost based on one or several villages with relatively small sample size, and serum samples from healthy volunteers were tested for SFTS virus-specific IgG or total antibodies (IgG and IgM) by using a double-antigen sandwich enzyme-linked immune sorbent assay (ELISA) kit or indirect-ELISA[24-27]. In past seroprevlence of SFTS virus investigations, the seropositive rate of healthy humans ranged from 0.23% to 9.17% varied in different regions[8]. The seroprevlence of SFTSV in Xinyang rural region was a little higher than that before. Because of small sample size or survey location in urban region or urban region, many seroprevlence of SFTS virus investigations may lead to lack of representativeness. However, in similar SFTS high endemic regions of Xinyang city, a seropositive rate of 6.59% was determined from 2011 to 2013[28]. The data showed that the seroprevlence of healthy humans was increasing compared to this study in 2016(10.46%), though there were some differences in testing or sampling method.

In this study, 12 healthy persons had positive SFTS virus-specific IgM antibodies. The results of follow-up showed that the healthy persons with IgM antibody were asymptomatic of SFTS, so they could be speculated as the SFTS virus covert infection. This study cannot rule out the possibility of these SFTS virus-specific IgM antibodies may have recall bias such as neglected mild clinical symptoms. Fortunately, 5 SFTS virus isolates were cultured from healthy individuals. The nucleotide sequences of SFTS virus isolates in this study were highly homologous, and had a high homology with viral nucleic acids and proteins isolated from SFTS patients, all belonging to the predominant epidemic A genotype in this area. The reason of covert infection is most likely that the virus loads in the body do not reach the pathogenic concentration, and the body depends on its own immune system to clear the virus without showing symptoms[29].

In Conclusion, the level of specific SFTS seroprevlence is relatively higher and possibility increasing in severe SFTS endemic region. There are asymptomatic SFTS case in healthy humans, so we suggested to develop vaccine research and take more improvements to ensure better prevention and control in high endemic region.

## Additional Information

### Publisher’s note

Springer Nature remains neutral with regard to jurisdictional claims in published maps and institutional affiliations.

